# Dissociable electrophysiological correlates of semantic access of motor and non-motor concepts

**DOI:** 10.1101/517284

**Authors:** Rodika Sokoliuk, Sara Calzolari, Damian Cruse

## Abstract

The notion of semantic embodiment posits that concepts are represented in the same neural sensorimotor systems that were involved in their acquisition. However, evidence in support of embodied semantics – in particular the hypothesised contribution of motor and premotor cortex to the representation of action concepts – is varied. Here, we tested the hypothesis that, consistent with semantic embodiment, sensorimotor cortices will rapidly become active while healthy participants access the meaning of visually-presented motor and non-motor action verbs. Event-related potentials revealed early differential processing of *motor* and *non-motor* verbs (164-203ms) within distinct regions of cortex likely reflecting rapid cortical activation of differentially distributed semantic representations. However, we found no evidence for a specific role of sensorimotor cortices in supporting these representations. Moreover, we observed a later modulation of the alpha band (8-12Hz) from 555-785ms over central electrodes, with estimated generators within the left superior parietal lobule, which may reflect post-lexical activation of the object-directed features of the motor action concepts. In conclusion, we find no evidence for a specific role of sensorimotor cortices when healthy participants judge the meaning of visually-presented action verbs. However, the relative contribution of sensorimotor cortices to action comprehension may vary as a function of task goals.

## Introduction

Theories of semantic embodiment propose that concepts are, at least in part, represented within the same neural sensorimotor systems that were involved in their acquisition (e.g. ^1,2^). Motor action concepts, for example, are considered to be represented within the brain’s motor cortices. In other words, the neuronal assemblies that represent the concept ‘to kick’ are thought to overlap with those involved in physically kicking one’s leg.

In support of the hypothesised overlap between action execution and action comprehension, there is considerable evidence that healthy participants are faster to perform actions in response to sentences if those actions are congruent with the actions described by the sentences – e.g. pushing a joystick away from you in response to the sentence “Close the drawer” ^3^. One study observed that healthy participants were selectively slower to make semantic judgments of sentences that describe actions if their motor system had been recently fatigued by repetitive action ^4^. Furthermore, there is evidence that participants experience interference in planning and executing hand actions when they are simultaneously required to access the meanings of visually presented action words ^5^. Functional neuroimaging data suggests that listening to sentences describing actions elicits greater activity in premotor cortex than listening to sentences that do not describe actions ^6^. Moreover, some studies have reported effector-specific overlap between tasks that involve action comprehension and action execution in premotor and primary motor cortices ^7–11^.

One commonly used marker of sensorimotor cortex activation is the modulation of electrophysiological oscillations in the mu (8-12Hz) and beta (13-30Hz) bands over the top of the head. Reductions in the amplitude of oscillations in these ranges in electroencephalography (EEG) and magnetoencephalography (MEG), often referred to as event related desynchronisations (ERD; ^12^), are ubiquitous when healthy individuals complete motor tasks, including imagery, observation, and execution (e.g. ^13–15^). These motor ERDs are often reported to be somatotopically distributed, suggesting sources within somatotopic cortices, although this is variable across studies ^16–20^. Nevertheless, data from simultaneous EEG-fMRI indicate broad somatosensory and motor cortical generators of the mu and beta rhythms in motor tasks^21^. Furthermore, the temporal resolution of electrophysiology allows researchers to separate cortical activity that may contribute to semantic access (i.e. within the first ~400 milliseconds post-stimulus;^22^) from that which occurs subsequent to semantic access, such as implicit or explicit mental imagery.

Consequently, and often cited in support of embodied semantics, somatotopy of MEG-recorded ERDs in the mu and beta bands have been reported from 200ms after presentation of action words^23^. Event-related potentials (ERPs) elicited by verbs and nouns also begin to differentiate from 200ms post-stimulus^24,25^ while source estimates of ERPs and MEG event-related fields have also suggested somatotopy of premotor and primary motor cortical contributions to semantic access of action words from 150ms post-stimulus^26–29^. The timing of this motor activation, hundreds of milliseconds before any overt semantic judgment by the participant, is viewed by some as support for the hypothesis that the motor system is engaged in service of semantic access of action concepts.

Nevertheless, post-lexical cortical activity has also been reported. For example, verbs and nouns have been associated with late (>500ms post-stimulus) topographically distinct oscillations in the beta range (25-35Hz)^24^. Larger beta ERDs have been observed in response to verbs relative to nouns from 600ms post-stimulus^30^ with putative generators in primary motor cortex^31^. However, the opposite pattern – i.e. larger ERDs for nouns relative to verbs over central scalp – has also been reported in the high beta / low gamma range^32^, suggesting the activity of multiple non-overlapping oscillatory mechanisms in semantic processing.

Perhaps mirroring the variability of the above evidence, a meta-analysis of fMRI and PET activation foci found insufficient evidence for the specific involvement of motor cortices in action verb processing, and instead observed a more consistent role for left lateral temporo-occipital cortex^33^. The authors concluded that action representations may overlap more with the cortical regions involved in perceiving actions (i.e. visual motion areas), rather than those involved in performing them (i.e. motor cortices). While this interpretation still falls within an embodied view of semantics, it calls into question the role of motor cortices in action verb comprehension. Nevertheless, there is evidence that motor cortical involvement in comprehension varies as a function of task goals^8,34^, and may therefore not be evident on average across a broad literature. Here we report an EEG study designed to investigate the specific cortical contributions to comprehension in the case of healthy participants accessing the meaning of visually-presented verbs.

Many high temporal resolution studies, reviewed above, have contrasted broad categories of words, such as verbs and nouns, which also differ on a range of potentially confounding psycholinguistic variables, such as imageability^35^. Furthermore, studies of neural responses to effector-specific words (e.g. lick, pick, kick) have relied on inverse models to accurately separate sub-regions of sensorimotor cortices^26,28,29^. Here, we describe a study of semantic access of verbs that differ in the extent to which they describe action but do not rely on source estimation of effector-specific sub-regions of motor cortices. This allows us to test the hypothesis that semantic access of motor verbs (e.g. ‘grab’) recruits dissociable regions of cortex from non-motor verbs (e.g. ‘fail’), as measured by ERPs and ERDs. By employing source estimation of these effects, we also tested whether the differential activity can be explained by overlap with cortical regions involved in performing actions (i.e. motor cortices) and those involved in perceiving actions (i.e. left lateral temporo-occipital cortex).

## Methods

### Participants

A total of forty-nine healthy participants (students from the University of Birmingham) took part in the studies and were compensated with either course credits or cash. Twenty of these participants took part in an initial behavioural study to validate the stimuli list (median age = 21.5; range: 18-31), and the remaining twenty-nine participants took part in the EEG study. Five participants were excluded from the EEG study due to excessive artefact, resulting in twenty-four participants (median age = 21; range: 19-28) for analysis. A sample size of 24 in a two-tailed within-subjects design gives 80% power to detect an effect size of .6^36^. All participants reported to be monolingual native English speakers, between 18 and 35 years old, right-handed, with no history of epilepsy, and no diagnosis of dyslexia. The experimental procedures were approved by the Ethical Review Committee of the University of Birmingham (ERN_15-1367AP3). All participants gave written informed consent.

### Stimuli

We constructed an initial list of 100 monosyllabic bodily action words and 100 monosyllabic non-bodily action words. We then used the *Match* software^37^ to select the stimuli set that best matched the motor and non-motor verbs on the basis of the following psycholinguistic variables: number of letters, number of phonemes, log frequency, orthographic neighbourhood, phonological neighbourhood, concreteness, imageability, and mean age of acquisition. The MRC Psycholinguistic Database^38^ provided the values for number of letters, number of phonemes, and concreteness. Word frequency values were taken from the British National Corpus Frequency database^39^. N-Watch software^40^ provided the orthographic and phonemic neighbourhood measures. Imageability ratings were taken from^41^, and age of acquisition ratings from^42^. This procedure resulted in a list of 36 motor and 36 non-motor verbs.

To validate this word list, we collected data from a group of healthy participants. Each participant was presented with each word individually, and instructed to create a sentence incorporating that word. Upon completion of this task, participants were again presented with each word and instructed to rate from 1 to 7 the extent to which the verb described a “voluntary and bodily action or movement” (7 as highest). We subsequently removed all words that were used as nouns by more than half of the participants or that received inconsistent ratings across participants (3 words per condition: tug, stroll, split, glare, glow, bleed). The resulting set of motor words were rated significantly higher than non-motor words (Wilcoxon’s Signed Rank Test, Z=561, p<.001), thus validating the motoric difference in meaning across lists while approximately controlling for potentially confounding psycholinguistic variables.

T-tests (Table 1) revealed no significant differences between the two final lists (33-words per condition) for any of the variables (*p*>.09). Bayesian T-tests (conducted with JASP v. 0.8. 0.0 software;^43,44^) revealed at least substantial evidence in favour of the null hypothesis for the majority of variables (i.e. BF_10_ ≤ 1/3), and weak evidence in favour of the null hypothesis for number of phonemes (BF_10_=.397), phonological neighbourhood (BF_10_=.600), and imageability (BF10=.831).

**Table 1.** Descriptive and inferential statistics for each psycholinguistic variable across Motor and Non-Motor stimuli. *For this comparison, we report Welch’s t-test for unequal variance as a Levene’s test indicated a violation of the assumption of equal variance.

|  | Motor |  | Non-motor |  | T-test |  |  | Bayesian T-test |
| --- | --- | --- | --- | --- | --- | --- | --- | --- |
|  | Mean | SD | Mean | SD | t | df | p | BF10 |
| **Log Frequency** | 1.004 | 0.636 | 0.967 | 0.524 | 0.257 | 64 | 0.798 | 0.259 |
| **Length in Letters** | 4.455 | 0.905 | 4.455 | 1.003 | 0.000 | 64 | 1.000 | 0.252 |
| **Length in Phonemes** | 3.727 | 0.761 | 3.545 | 0.666 | 1.033 | 64 | 0.306 | 0.397 |
| **Orthographic Neighbourhood** | 5.636 | 4.076 | 5.879 | 4.121 | -0.240 | 64 | 0.811 | 0.258 |
| **Phonological Neighbourhood** | 12.818 | 6.361 | 15.455 | 8.449 | -1.432 | 64 | 0.157 | 0.600 |
| **Imageability\*** | 4.194 | 0.499 | 3.924 | 0.773 | 1.683 | 54.7 | 0.098 | 0.831 |
| **Mean Age of Acquisition** | 6.205 | 1.906 | 6.368 | 1.695 | -0.367 | 64 | 0.715 | 0.267 |

### Procedure

The paradigm was programmed and presented using the Matlab Psychophysics Toolbox (Matlab Psychtoolbox-3; www.psychtoolbox.org^45^). Participants sat approximately 100 cm from a 27-inch PC monitor, with refresh rate of 60 Hz, 1920×1080 resolution, and 32-bit colour depth.

Each trial (Figure 1) began with a central grey fixation point on a black background for 1500-ms, followed by a central white fixation cross for 200-ms, a blank screen for 1300-ms, and finally the word presented in lower case Arial (size 80) at the centre of the screen for 200-ms, followed by 1300-ms of blank screen. To promote participant attention to the meaning of the stimuli, 25% of trials were followed by presentation of a word definition, taken from web-based dictionaries, to which the participant was required to judge whether the definition matched the preceding word. Responses were given via keyboard, with response hand counterbalanced across participants (i.e. left-hand to answer “correct” and right-hand to answer “incorrect” for half of the participants, and vice versa for the other half). Definitions matched the preceding word exactly half of the time. Stimulus order and the stimuli chosen for presentation of definitions were randomised. Due to a bug in the presentation script, the order of stimuli was identical for half of the participants. Nevertheless, the order of stimuli for those participants was unpredictable. At the end of every trial, a blank screen was presented for between 1000- and 2000-ms, selected on each trial from a uniform distribution.

**Figure 1:**
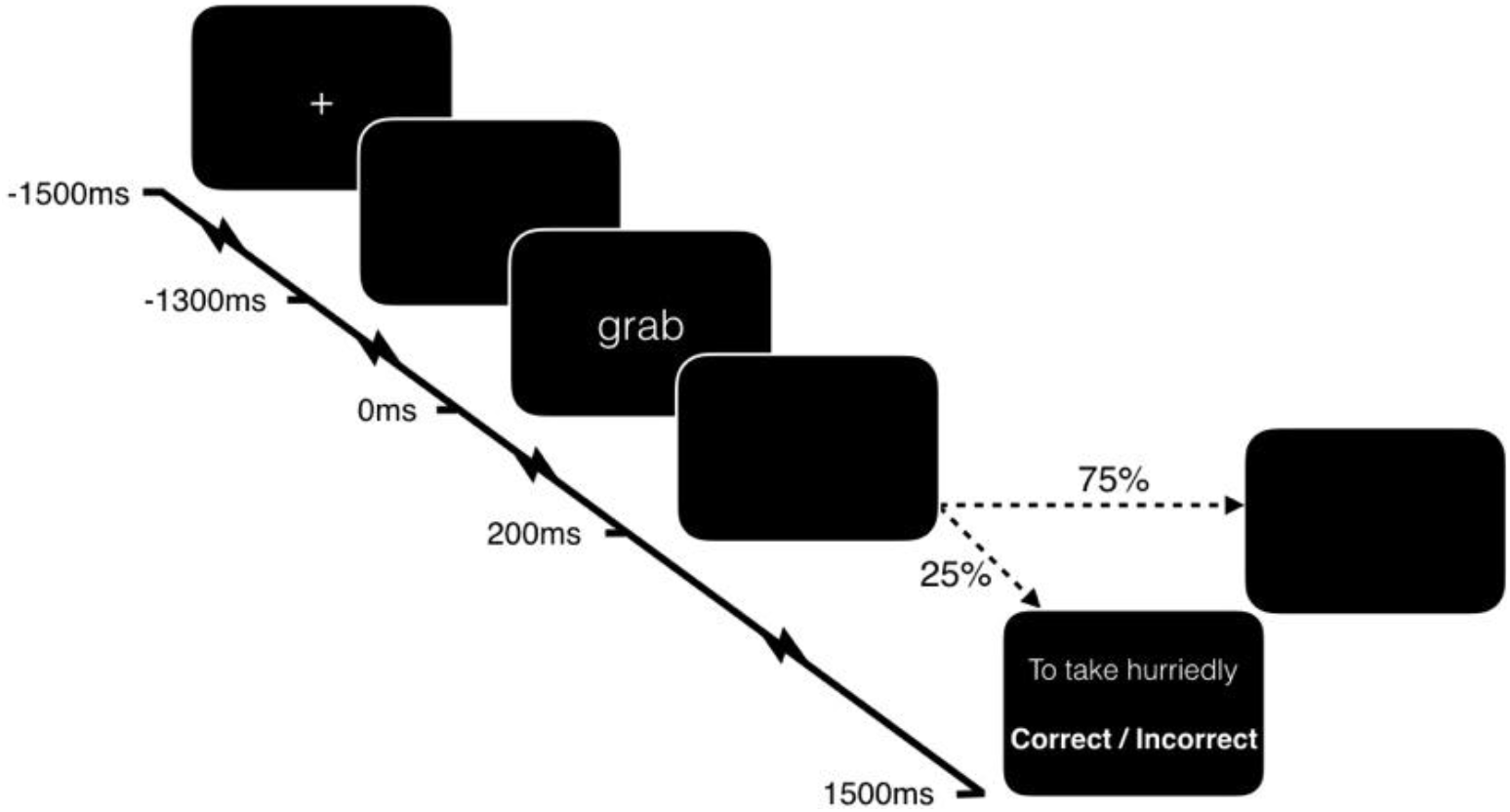
Trial procedure with timings relative to stimulus presentation

To improve signal to noise, participants completed four runs of the above procedure, resulting in 132 trials per condition. Across all 4 runs, a definition for each word was presented exactly once. Participants also completed a brief practice session of six trials to familiarise to the structure of the task. Practice stimuli were the words rejected during the stimuli validation procedure described in the *Stimuli* section above.

### EEG pre-processing

We recorded EEG with a 128-channel Biosemi ActiveTwo system, with two additional electrodes recording from the mastoid processes. Data were sampled at 256 Hz and referenced to CMS (Common Mode Sense) and DRL (Driven Right Leg). Offline, the EEG signal was digitally filtered between 0.5 and 40 Hz, segmented into epochs from 1500-ms prestimulus until 1500-ms poststimulus, re-referenced to the average of the mastoids, and baseline-corrected to the 200-ms prestimulus period. All offline pre-processing was performed with a combination of the Matlab toolbox EEGLAB (version 14.0.0b^46^) and custom scripts.

Artefact rejection proceeded in three steps. First, channels and trials with excessive or non-stationary artefact were identified by visual inspection and discarded. Across participants, a median of 4 channels (range 0-12) and a median of 38.5 trials (range 11-73) were discarded. Second, we conducted Independent Component Analysis (ICA) of the remaining data (EEGLAB’s *runica* algorithm) to identify and remove components that described eye blinks and eye movements. Any previously removed channels were then interpolated back into the data. Finally, trials with artefacts that had not been effectively cleaned by the above procedure were identified with visual inspection and discarded.

Prior to analysis, all data were re-referenced to the average of all channels, and baseline corrected to the 200-ms prestimulus period. A median of 113.5 trials per participant contributed to each condition (Motor range: 93-127; Non-motor range: 98-127).

### EEG / MRI co-registration

We recorded the electrode positions of each participant relative to the surface of the head with a Polhemus Fastrak device using the Brainstorm Digitize application (Brainstorm v. 3.4^47^) running in Matlab. Furthermore, on a separate day, we acquired a T_1_-weighted anatomical scan of the head (nose included) of each participant with a 1mm resolution using a 3T Philips Achieva MRI scanner (32 channel head coil). This T_1_-weighted anatomical scan was then co-registered with the digitised electrode locations using Fieldtrip^48^.

### Sensor analyses: ERPs

Analyses of ERPs proceeded in two stages. First, we calculated the global field power^49^ of the grand average of all trials (i.e. both conditions together) to identify time-windows of interest. Global field power (GFP) is the root mean square of average-referenced voltages, and is a principled means of identifying component peak latencies from an orthogonal contrast. We then identified a time-window around each peak by inspecting the global dissimilarity^49^ – the mean of the root mean square of voltage differences between consecutive time-points, after the data have been scaled by the global field power. Deflections in the time-course of global dissimilarity therefore suggest boundaries between scalp topographies. Due to the focus on processing in support of semantic access, we selected four ERP topographies approximately within the first 400-ms post-stimulus: 106-ms – 160-ms, 164-ms – 203-ms, 207-ms – 293-ms, and 297-ms – 418-ms (see Figure 2).

**Figure 2:**
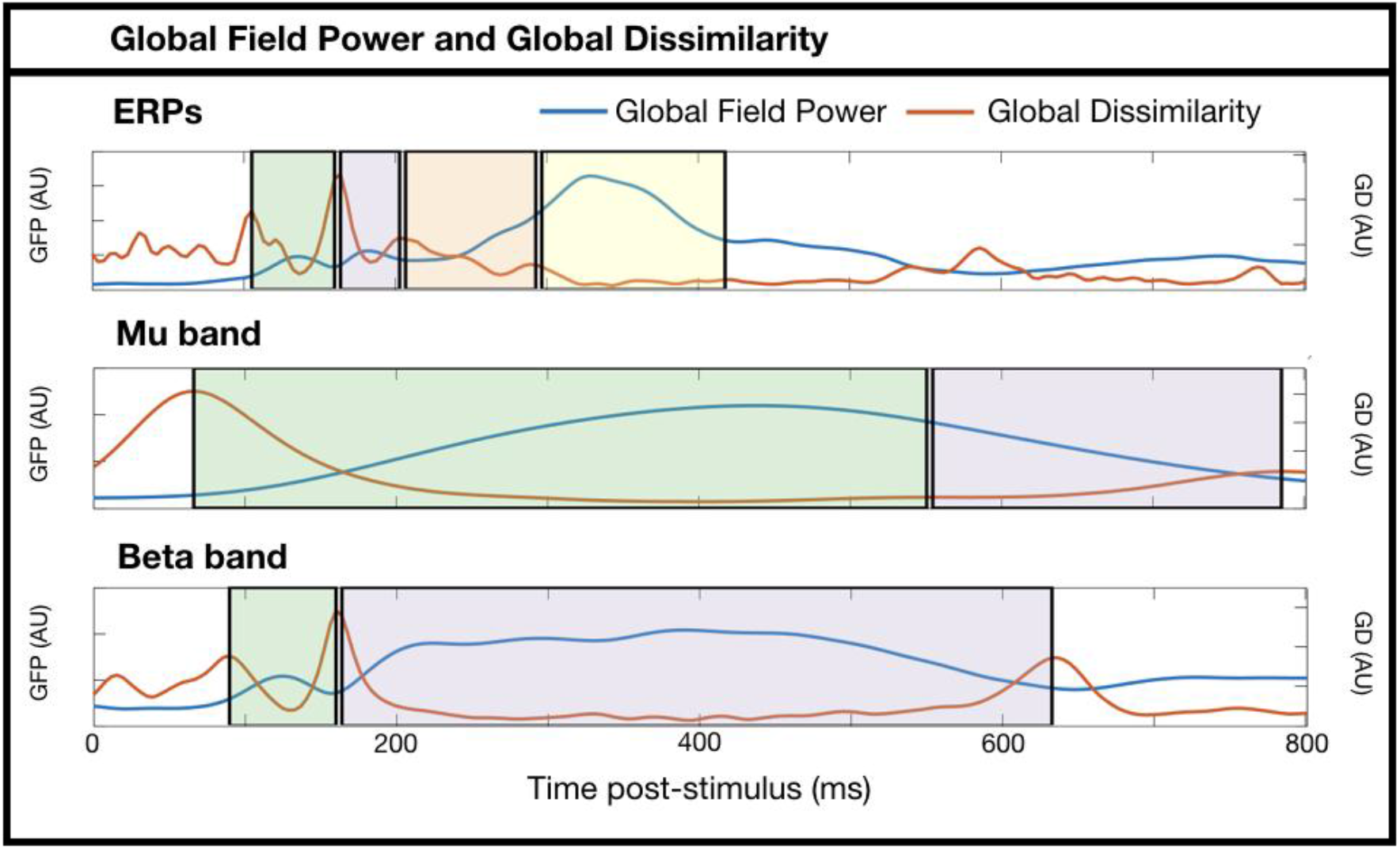
Global Field Power (GFP) and Global Dissimilarity (GD) time-courses of the ERPs and mu and beta band-power estimates. Shaded areas represent the time-windows selected for subsequent analyses. Note that these are plots of grand average data across conditions, and are therefore orthogonal to the subsequent motor versus non-motor analyses.

ERPs within each time-window of interest were compared with the cluster mass method of the open-source Matlab toolbox FieldTrip (version 20160619,^48^). First, for each participant x condition we averaged the voltages at each electrode within the time-window of interest. Next, a two-tailed dependent samples t-test between conditions was conducted at each electrode. Spatially adjacent t-values with p-values passing the threshold (alpha = .05) were then clustered based on their spatial proximity. Spatial clusters were required to involve at least 4 neighbouring electrodes. To correct for multiple comparisons, a randomisation procedure produced 1000 Monte Carlo permutations of the above method to estimate the probability of the observed cluster under the null hypothesis^50^. We used a cluster alpha threshold of .025 as we are testing for both positive and negative effects.

As we hypothesise that the neural representations of motor and non-motor verbs are not entirely overlapping, we also tested for differences in the scalp topographies across conditions with a randomisation test of global dissimilarity (see^51^). For each time-window of interest, we calculated the global dissimilarity (i.e. the root mean square difference in GFP-normalised voltages) between the grand-average topographies of the two conditions. We then estimated the probability of observing that global dissimilarity (or a value larger) under the null hypothesis. Specifically, we randomly shuffled data across conditions, while maintaining within-subject pairings of condition, and re-calculated global dissimilarity as above. The p-value is the proportion of global dissimilarities from 1000 randomisations that are larger than the observed global dissimilarity.

### Sensor analyses: oscillations

To estimate power in each frequency band of interest (mu: 8-12Hz; beta: 13-30Hz) we filtered all individual trials within the band of interest (EEGLAB *firls*) and extracted the squared envelope of the signal (i.e. the squared complex magnitude of the Hilbert-transformed signal).

We then averaged trials of the same condition within each participant’s data, and converted post-stimulus values to decibels relative to the mean power in a pre-stimulus baseline (−600 to −200ms) selected to not be contaminated by temporally-smeared post-stimulus power estimates. Subsequent statistical procedures were identical to the ERP sensor analyses above. Due to previous evidence of late oscillatory changes during verb processing, we identified time-windows within the first 800-ms post-stimulus from the GFP and GD time-courses of the two frequency bands: mu: 66-ms – 551-ms, 555-ms – 785-ms; beta: 90-ms – 160-ms, 164-ms – 633-ms (see Figure 2).

### Source analysis

We performed source analyses on data of 20 out of the 24 participants because we were unable to acquire anatomical MRI scans for the remaining four participants. From the subject-specific T_1_-weighted anatomical scans, individual boundary element head models (BEM; four layers) were constructed using the ‘dipoli’ method of the Matlab toolbox FieldTrip^48^. Digitised electrode locations were aligned to the surface of the scalp layer that was extracted from the segmented T_1_-weighted anatomical scans using fiducial points and head shape as reference points.

Data that was analysed previously on the sensor level was now projected onto the source level. To allow for direct statistical comparison between *motor* vs. *non-motor* verbs, the number of trials was balanced between conditions by randomly removing trials of the condition (discarded trials: median 3, range 0-10) with more data until both datasets had the same number of trials (median 112, range 94-125).

#### ERPs whole brain

For the ERP source dataset, trials were defined as time windows reaching from [-400ms – 1200ms]. To increase computational efficiency these individual trials were concatenated over time for each subject separately. Using a Linear Constraint Minimum Variance (LCMV) beamformer^52–54^, a spatial filter was constructed. To obtain an estimate of the overall brain response to *motor* vs. *non-motor* verbs, all virtual electrodes (VEs) were extracted and their time course computed. The continuous time courses were then epoched into individual trials of 1.2s [-200ms – 1000ms], excluding the first and last 200ms of the original trial length to avoid including potential artefacts due to data discontinuities. These data were then separated into *motor* and *non-motor* trials and averaged over trials for each condition separately. Then these data were baseline-corrected using the time window [-200ms – 0ms] relative to stimulus presentation and further normalized by the standard deviation over the baseline period to reduce inter-subject variability. Data were further averaged over the time window that showed significant clusters on the sensor level (164-203ms) before the difference between *motor* and *non-motor* condition was computed for every participant. The overall brain response, averaged over 20 participants, is represented in figure 6.

#### Mu oscillations whole brain

For the mu oscillation dataset, trials were defined as time windows reaching from [-800ms – 1200ms] relative to stimulus presentation. To increase computational efficiency, individual trials were concatenated, resulting in one continuous datastream for each subject. A Linear Constraint Minimum Variance (LCMV) beamformer^52–54^ was used to construct a spatial filter. Then, all VEs were extracted and their time course computed. To obtain mu power values, time courses were further hilbert-transformed and their absolute values squared. Data were then epoched into individual trials of 1.6s [-600ms – 1000ms], excluding the first and last 200ms to avoid potential artefacts due to data discontinuities. Then, trials were split into *motor* and *non-motor* conditions and the average over trials within each condition was computed. After baseline-correcting the data to dB using the pre-stimulus time window [−600ms to −200ms], average values over the time window that showed a significant cluster on the sensor level between *motor* and *non-motor* conditions (555ms – 785ms) were computed. Figure 7A shows the overall brain response to the difference in mu power between *motor* and *non-motor* verbs, averaged over 20 participants.

#### Automated Anatomical Labelling (AAL) analysis

We focused our analyses on the specific anatomical regions identified in the meta-analysis of Watson et al. (2013)^33^ described in the introduction to this manuscript, and bilateral precentral gyri based on evidence from literature^7–11^. The resulting seven anatomical regions of interest (see Table 2 and Figure 3 for anatomical details) were defined using the automated anatomical labelling (AAL) atlas (see^55,56^ for similar analyses with MEG and EEG data). One of these seven AAL regions, the left middle temporal gyrus (left MTG) was further subdivided into anterior (aMTG) and posterior middle temporal gyrus (pMTG) since only the left pMTG was of interest in this study (cf.^33^). This was done by selecting only those left MTG VEs located posterior to the centre of mass of the left MTG AAL region.

**Figure 3:**
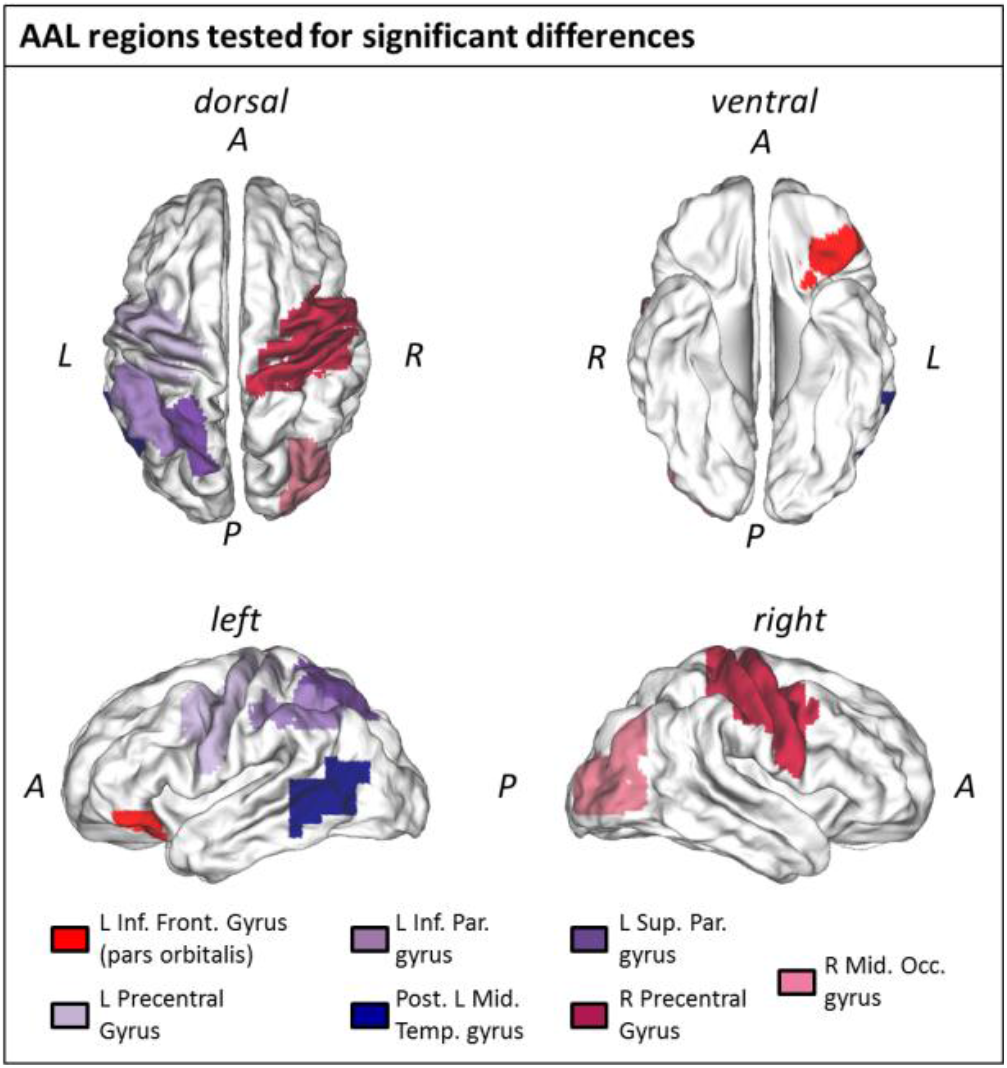
Locations of the seven investigated AAL regions (A = anterior; P = posterior; L = left; R = right).

**Table 2.** MNI coordinates of the centre of mass of all investigated AAL regions and peak locations from the associated meta-analysis study (Watson et al, 2013^33^). White fields in the centre column indicate MNI x, y, and z coordinates of AAL regions selected based on peak locations of a meta-analysis study of PET and fMRI action vs. non-action contrasts by Watson et al., 2013, shown in the right column. Grey fields represent MNI coordinates of two additional AAL regions investigated based on prior evidence showing that these regions are involved in action word processing (see Introduction).

| AAL region | AAL Centre of mass<br>MNI-coordinates [cm] |  |  | Peak meta-analysis<br>MNI-coordinates [cm] |  |  |
| --- | --- | --- | --- | --- | --- | --- |
|  | *x* | *y* | *z* | *x* | *y* | *z* |
| *R Occipital Mid. Gyrus* | 3.5 | -7.5 | 2 | 5.4 | -7 | 0.2 |
| *L Sup. Parietal Lobule* | -2.5 | -6 | 5.5 | -3.2 | -5.6 | 5.4 |
| *L Inf. Parietal Lobule* | -4 | -4.5 | 4.5 | -3.2 | -5 | 5.4 |
| *L Inf. Frontal Gyrus* | -3 | 2.5 | -1.5 | -4.4 | 3.2 | -1.6 |
| *L Post. Middle Temporal Gyrus* | -5 | -5.5 | -0.5 | -5.8 | -5 | 0.6 |
| *L Pre-central Gyrus* | -3.5 | -1 | 4.5 |  |  |  |
| *R Pre-central Gyrus* | 4 | -1 | 5 |  |  |  |

In a first step, time courses of all AAL regions’ VEs were extracted and weighted, based on the Euclidian distance between each VE and the centre of mass of the respective AAL region (cf.^55^). In order to investigate ERP results, time courses were summed across VEs within each AAL region separately and further processed for potential ERP source differences as described above (*ERPs whole brain*). To extract potential mu power differences on the source level between the conditions *motor* vs. *non-motor*, the time courses of all VEs were first Hilbert transformed and the absolute values squared before summing across VEs. Further processing followed the procedure described above (*Mu oscillations whole brain*). To test for statistical significance, paired-sample t-tests were performed on both datasets on the computed differences between *motor* and *non-motor* conditions for ERPs and mu oscillations (20 vs. 20 for each AAL region). Resulting p-values were further corrected for multiple comparisons using False Discovery Rate (FDR;^57,58^). For the AAL region showing significant differences between *motor* and *non-motor* conditions, additional one-sample t-tests were computed in order to test whether the mean of their distributions differed significantly from 0.

To test for evidence for the Null, Bayes Factor analyses with default priors (r=0.707) were carried out on the ERP and Mu data for each AAL region separately according to^59^.

### Plotting

All data plots were made with Matlab and edited in a desktop publisher. Colour palettes are taken from Color Brewer 2 (http://colorbrewer2.org/) or in-house customized colour maps. Source results were plotted using caret (http://www.nitrc.org/projects/caret/;^60^).

### Data Availability

The datasets generated during and/or analysed during the current study, along with all processing and analysis scripts, are available in the *Data* (https://osf.io/thznk/?view_only=f9302d298aca40bbb2f17b3c7291b981) and *Analysis Scripts* (https://osf.io/rwvxh/?view_only=af640c21223f41068af732667da474af) repositories.

## Results

### Behaviour

Participants judged the correctness of verb definitions with high accuracy (hit rate: M=91.79%, SD=11.25%; false alarm rate: M=3.03%, SD=2.96%) which we interpret as evidence of the group’s attention to the meaning of stimuli.

### Sensor analyses: ERPs

One positive and one negative spatial cluster exceeded our significance threshold in the 164-203-ms time-window (p=.008 and p=.002, respectively). The two clusters are located over both sides of a dipolar distribution of voltage differences, with an anterior positivity and posterior negativity (Figure 4B). Global dissimilarity within the 164-203-ms time-window was significantly greater than that expected by chance (GD=.416, p=.004), suggesting that the neural generators underlying motor and non-motor verb processing are not entirely overlapping in this time-window.

**Figure 4:**
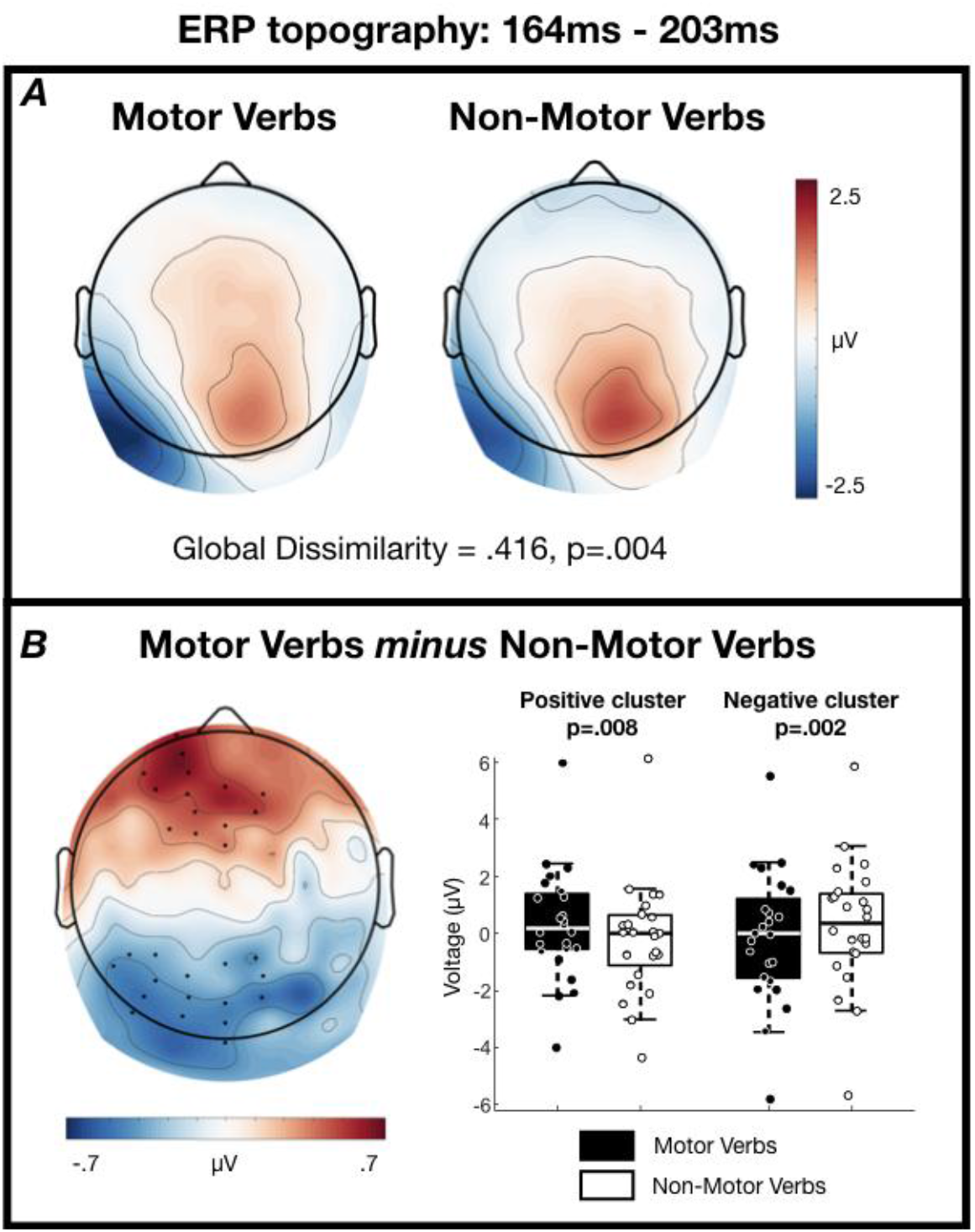
ERP scalp topographies from 164-203ms post-stimulus. (A) Grand average topographies separated according to condition. (B, left) Grand average topography of the difference between conditions. Electrodes contributing to the two clusters are marked. (B, right) Tukey boxplots and individual subject mean voltages within the two significant clusters.

No clusters were formed in any of the other three time-windows of interest, nor did any other global dissimilarity analyses exceed our statistical threshold (106-160-ms: GD .160, p=.998; 207-293-ms: GD .148, p=.325; 297-418-ms: GD .066, p=.399).

### Sensor analyses: Oscillations

Global dissimilarity within the 555-785-ms time-window of the mu band was significantly greater than that expected by chance (GD=.385, p=.009; Figure 5A), indicating that the neural generators underlying the mu band reactivity to motor and non-motor verbs are not entirely overlapping in this time-window. The scalp maps in Figure 5A clearly show occipital alpha reactivity in this time-window, with a greater extent of the motor verb ERD over central midline electrodes. In the same time-window, one cluster of voltage differences was formed with p=.050 centred on electrode Cz, although this fails to pass our two-tailed threshold of p<.025.

**Figure 5:**
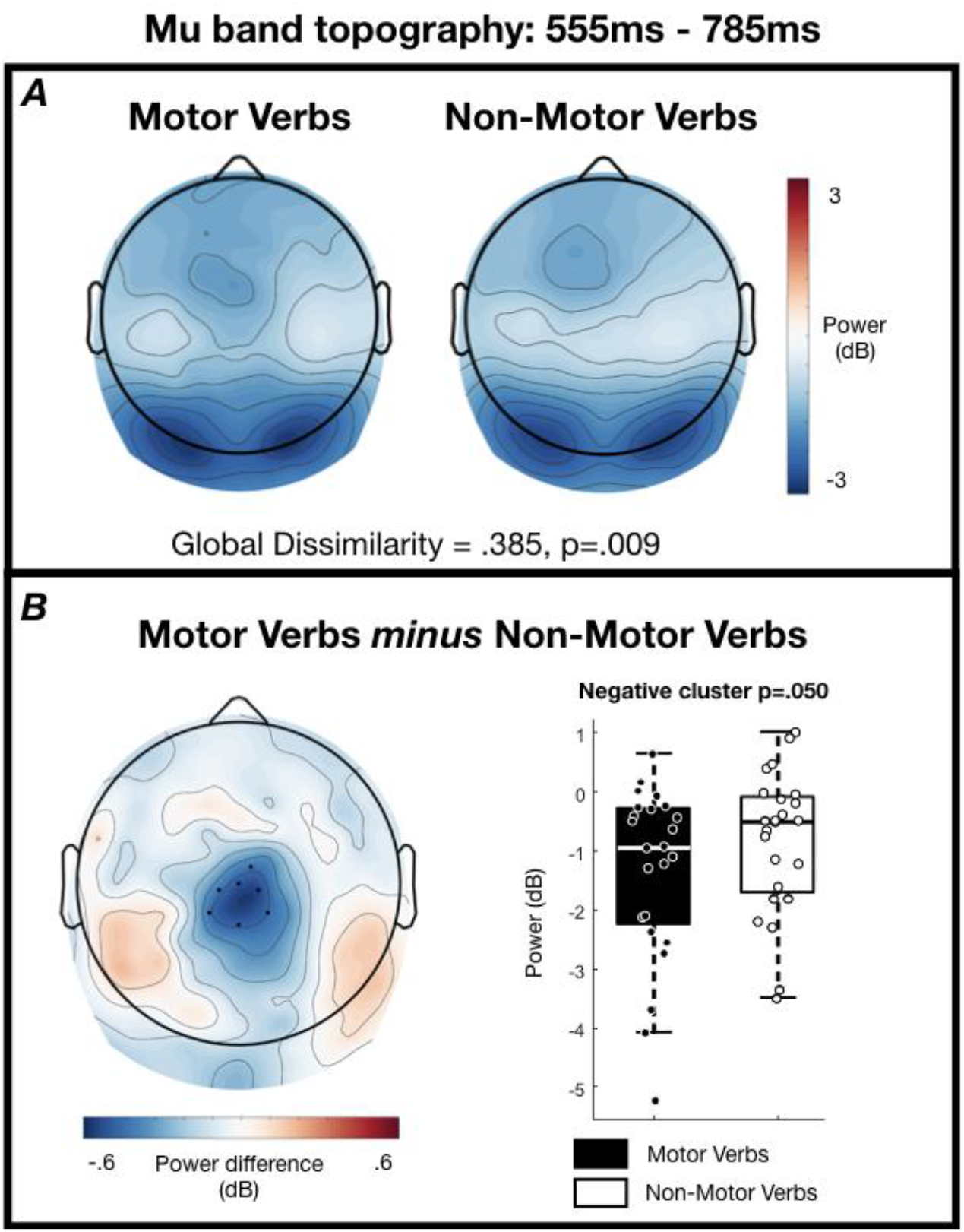
Mu band scalp topographies from 555-785ms post-stimulus. (A) Grand average topographies separated according to condition. (B, left) Grand average topography of the difference between conditions. Electrodes contributing to the cluster are marked. Note that this cluster does not pass the threshold of p<.025, but is shown to visualise the difference that drives the significant global dissimilarity (p=.009). (B, right) Tukey boxplots and individual subject mean power within the cluster of electrodes.

No clusters were formed in the early time-window for the mu band (66-ms – 551-ms), or in either of the time-windows for the beta band. No other global dissimilarity analyses exceeded our statistical threshold (Mu band 66-551-ms: GD .216, p=.143; Beta band 90-160-ms: GD .769, p=.364; Beta band 164-633-ms: GD.240, p=.685).

### Source analyses: ERP

Source analyses on the ERP dataset did not reveal any significant differences between *motor* and *non-motor* verbs in any of the tested AAL regions (paired one-sample t-test: FDR corrected p-values >0.05). Figure 6 shows difference values between ERP to *motor* and *non-motor* verbs averaged over participants and the time window that showed significant clusters on the sensor level (164-203ms). To test whether there are AAL regions that respond in the same way to motor and non-motor verbs within the analysed time window, a Bayesian t test was computed for each AAL region. It revealed some evidence for the Null in the left inferior parietal lobule (BF_10_ = 0.233; see Table 3 for detailed description of statistical tests), suggesting that the signal in the analysed time window of the ERP in this region is the same for motor and non-motor verbs.

**Figure 6:**
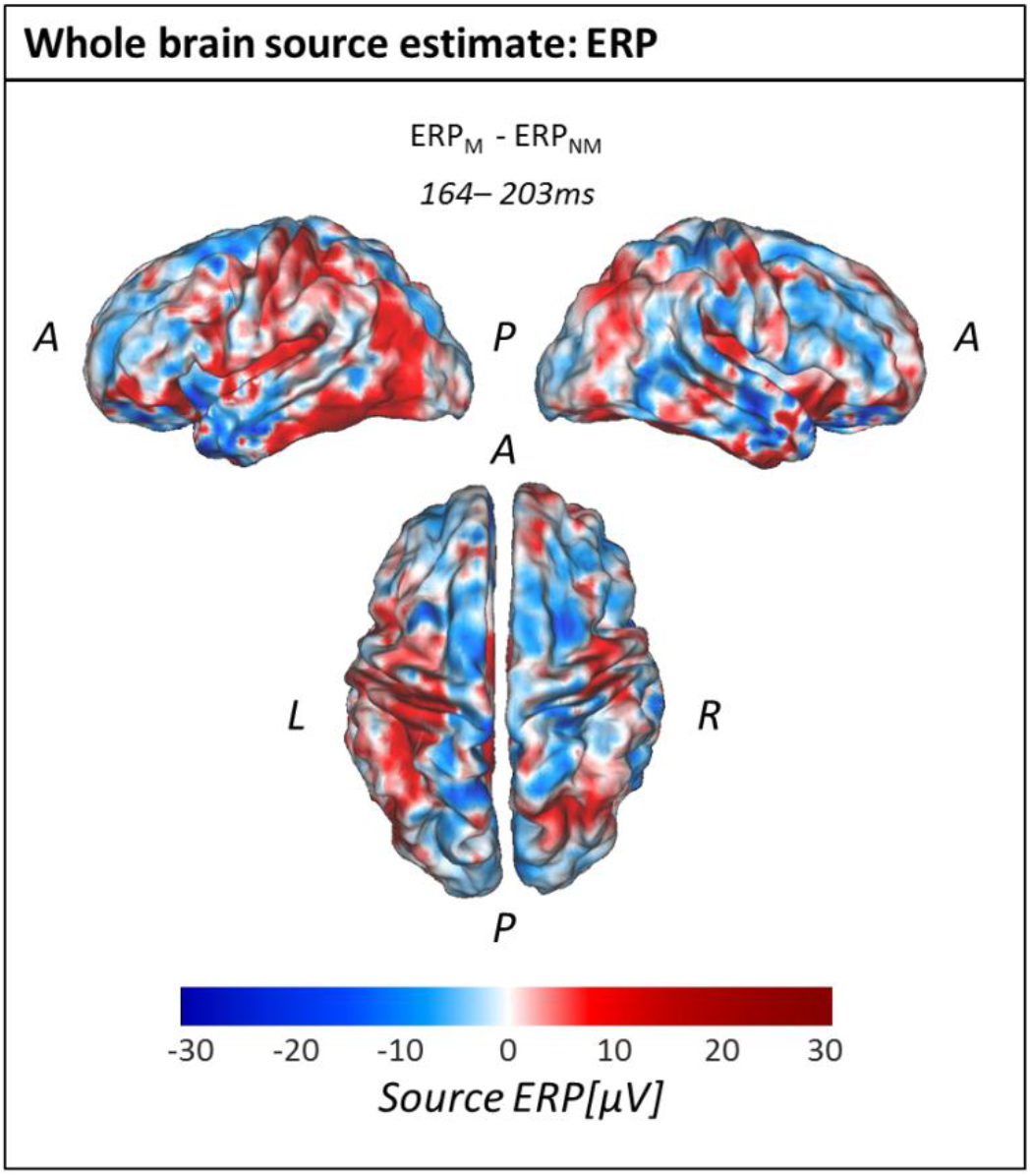
Whole brain source estimate for the difference between ERPs to motor and non-motor verbs averaged across the time-window of significant difference in the sensor data (164-203ms) [A = anterior; P = posterior; L = left; R = right].

**Table 3.**
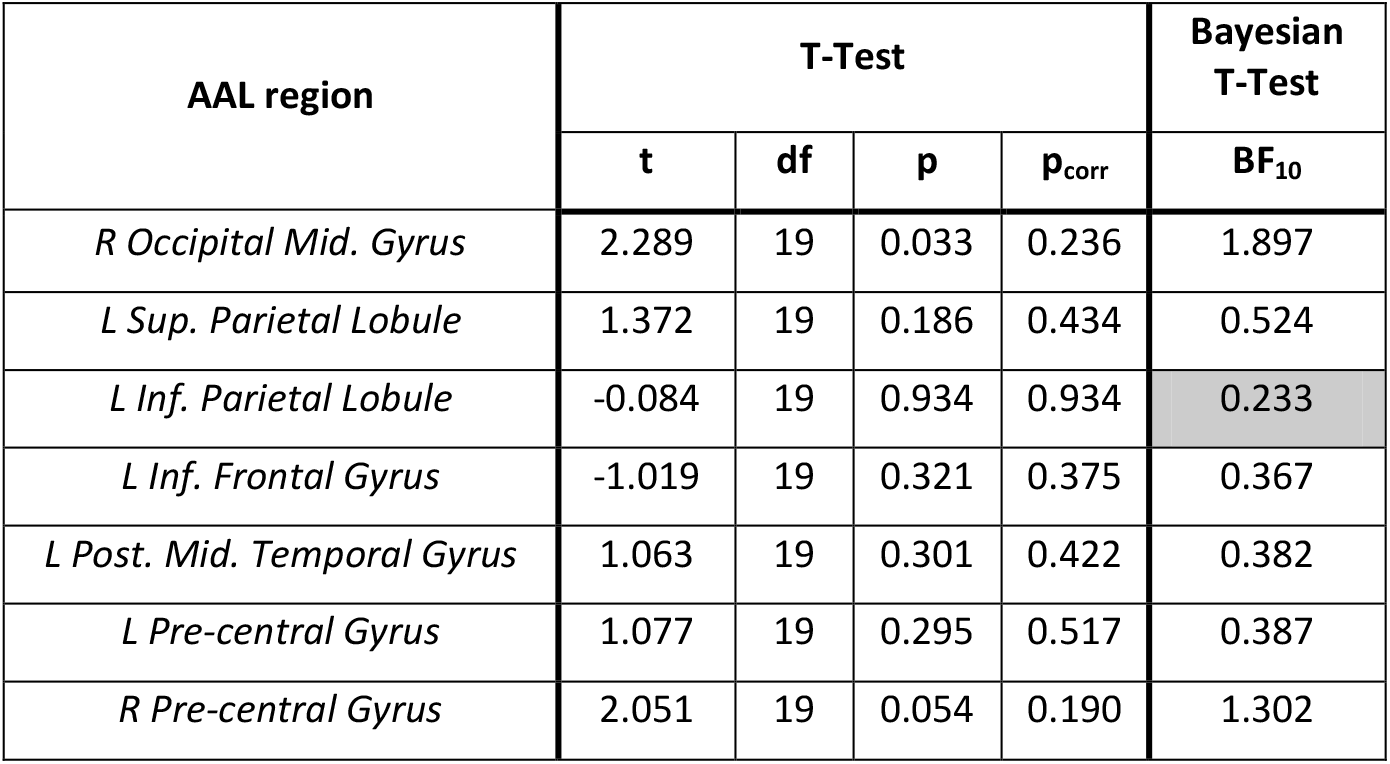
Paired sample T-Tests and Bayesian T-Tests of ERP source estimate in individual AAL regions. For every analysed AAL region, the T-value (t), the degree of freedom (df), the p-value (p), the FDR-corrected p-value and the Bayes factor (BF_10_ = support for H1 over H0; BF_10_<0.333: substantial evidence for the Null) are presented. Grey shaded Bayes factor represents substantial evidence for the Null in the left Inferior Parietal Lobule. “R” and “L” in left column identify AAL regions in the “right” and “left” hemisphere respectively.

| AAL region | T-Test |  |  |  | Bayesian T-Test |
| --- | --- | --- | --- | --- | --- |
|  | t | df | p | p<sub>corr</sub> | BF<sub>10</sub> |
| *R Occipital Mid. Gyrus* | 2.289 | 19 | 0.033 | 0.236 | 1.897 |
| *L Sup. Parietal Lobule* | 1.372 | 19 | 0.186 | 0.434 | 0.524 |
| *L Inf. Parietal Lobule* | -0.084 | 19 | 0.934 | 0.934 | 0.233 |
| *L Inf. Frontal Gyrus* | -1.019 | 19 | 0.321 | 0.375 | 0.367 |
| *L Post. Mid. Temporal Gyrus* | 1.063 | 19 | 0.301 | 0.422 | 0.382 |
| *L Pre-central Gyrus* | 1.077 | 19 | 0.295 | 0.517 | 0.387 |
| *R Pre-central Gyrus* | 2.051 | 19 | 0.054 | 0.190 | 1.302 |

### Source analyses: Mu/alpha-oscillations

Source estimates of the differential mu/alpha (see Discussion for consideration of whether this is a mu or alpha rhythm) response during the time-window of the significant effect at the sensor level reveal a broad negativity over centro-posterior brain regions (see figure 7A). The difference in the left superior parietal lobule survived multiple comparisons correction, reflecting a reduction in mu/alpha power in response to motor verbs relative to an increase in mu/alpha power to non-motor verbs (T(19) = −3.249; p = 0.029 FDR corrected; figure 7B), which was further confirmed by a Bayesian t test (BF_10_ = 10.598). Mu/alpha power distributions of *motor* and *non-motor* conditions in that AAL region however were not significantly different from 0 (*motor*: T(19) = 0.107; *non-motor:* T(19) = 0.604). Applying the Bayesian t test to all other AAL regions (see Table 4 for detailed description of statistical tests) showed some evidence for the Null in the posterior middle temporal gyrus (BF_10_ = 0.232), the left pre-central gyrus (BF_10_ = 0.237), and the right middle occipital gyrus (BF_10_ = 0.251), suggesting that mu/alpha power in these regions is not modulated differently upon processing of motor or non-motor words.

**Figure 7:**
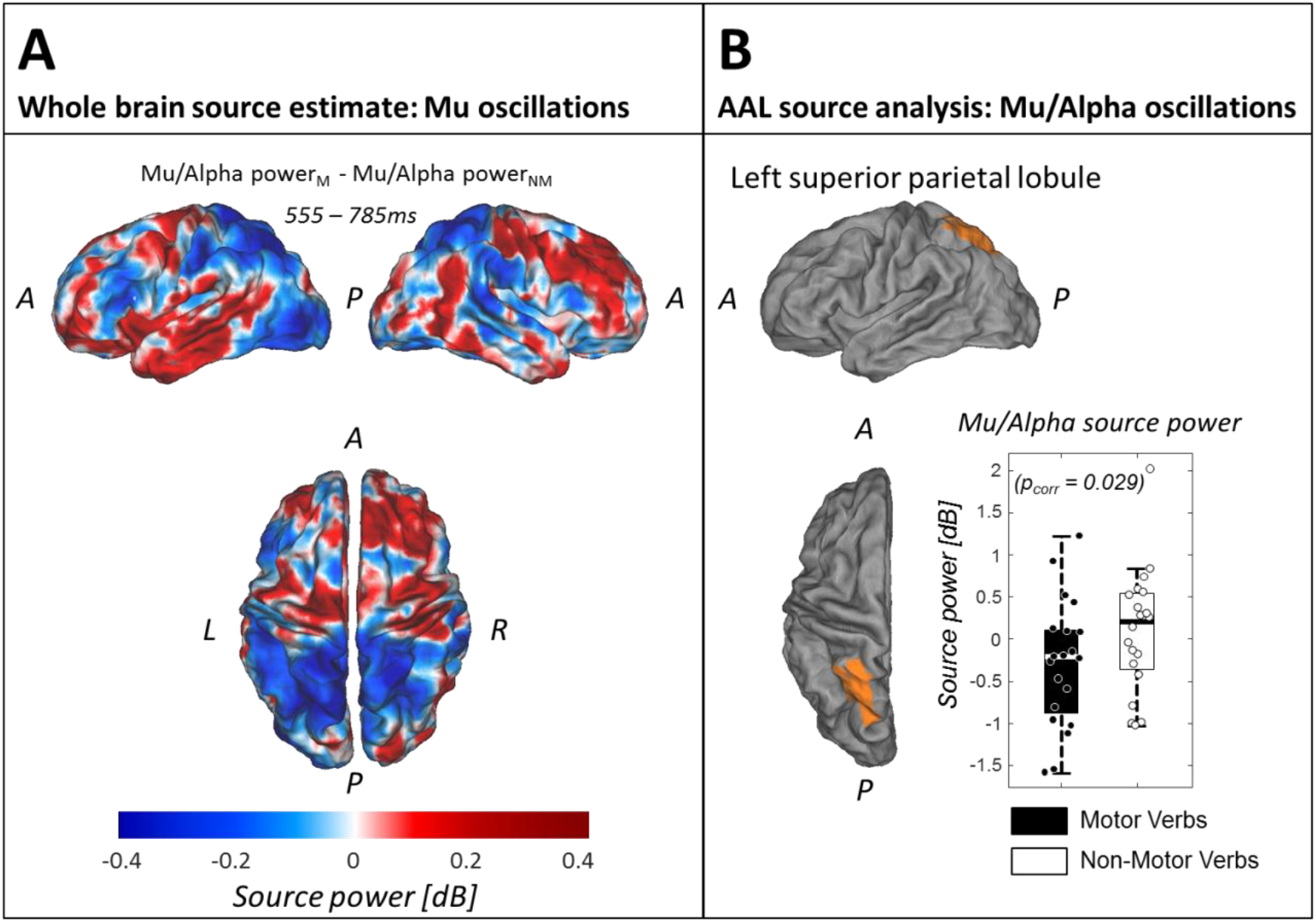
Source results Mu/Alpha oscillations. (A) Mu/alpha power difference values averaged over participants and the time window that showed a significant negative cluster on the sensor level (555-785ms). (B) AAL region ‘left superior parietal lobule’ (in orange) that showed a significant mu/alpha power difference between motor and non-motor conditions (p = 0.029; FDR-corrected). The Tukey boxplots represent mu/alpha power values in the AAL region ‘left superior parietal lobule’ averaged over the time window of interest (555-785ms) for motor and non-motor conditions, with individual subject means overlaid. Abbreviations identify spatial landmarks: A = anterior; P = posterior; L = left; R = right.)

**Table 4.** Paired sample T-Tests and Bayesian T-Tests of mu/alpha power source estimate in individual AAL regions. For every analysed AAL region, the T-value (t), the degree of freedom (df), the p-value (p), the FDR-corrected p-value and the Bayes factor (BF_10_ = support for H1 over H0; BF_10_<0.333: substantial evidence for the Null) are presented. Light grey shaded corrected p-value and Bayes Factor represent significant difference in mu/alpha power between motor and non-motor conditions. Dark grey shaded Bayes factors represent substantial evidence for the Null in three AAL regions. “R” and “L” in left column identify AAL regions in the “right” and “left” hemisphere respectively.

| AAL region | T-Test |  |  |  | Bayesian T-Test |
| --- | --- | --- | --- | --- | --- |
|  | t | df | p | p<sub>corr</sub> | BF<sub>10</sub> |
| *R Occipital Mid. Gyrus* | -0.412 | 19 | 0.685 | 0.959 | 0.251 |
| *L Sup. Parietal Lobule* | -3.251 | 19 | 0.004 | 0.029 | 10.598 |
| *L Inf. Parietal Lobule* | -2.115 | 19 | 0.048 | 0.168 | 1.438 |
| *L Inf. Frontal Gyrus* | 1.265 | 19 | 0.221 | 0.516 | 0.466 |
| *L Post. Mid. Temporal Gyrus* | 0.045 | 19 | 0.965 | 0.965 | 0.232 |
| *L Pre-central Gyrus* | 0.198 | 19 | 0.845 | 0.986 | 0.237 |
| *R Pre-central Gyrus* | 1.077 | 19 | 0.295 | 0.516 | 0.387 |

## Discussion

We tested the hypothesis that semantic access of verbs that describe motor actions involves dissociable regions of cortex from semantic access of verbs that do not describe motor actions. In support of this hypothesis, we observed significant differences between the ERPs elicited by these two classes of stimuli in a time-window 164-203ms post-stimulus. Global dissimilarity analysis in this time-window provided strong evidence that the ERPs in response to motor and non-motor verbs are not generated by entirely overlapping regions of cortex (p=.004). This result is consistent with widely-accepted distributed views in which a concept’s semantic features are represented across cortices (see, for example,^61^), but is not in itself sufficiently consistent with an embodied view of semantics, in which action concepts are considered to be represented within the cortical sensorimotor system.

The timing of our observed ERP effect is consistent with that reported in a study of Dutch arm action verbs and non-action verbs, in which ERPs diverged from 155-174ms post-stimulus^35^. Furthermore, source estimation implicated bilateral motor cortices (precentral gyri) as generators of that effect, and was therefore interpreted as evidence for embodied semantics of action. While the early onset of our observed ERP difference is consistent with cortical activity in support of semantic access^22^, we found no evidence that this ERP effect originated within cortical regions that would be consistent with embodied semantics – e.g. occipito-temporal (perceptual) areas identified in a meta-analysis of fMRI and PET results^33^ or bilateral pre- and primary motor cortices (see Table 2 and 3). Indeed, our Bayesian analyses suggested substantial evidence for the null hypothesis of no effect in left inferior parietal lobule (BF_10_=0.233) and weak evidence for the null in other linguistic and motor regions, including left pre-central gyrus (BF_10_=0.387).

One possible reason for the conflicting source estimates between our study and that of Vanhoutte et al.^35^ is that the verbs they used differed considerably in their imageability ratings between action and non-action categories (reported p<.001) while the verbs in our study did not differ significantly between categories (p=.098) and a Bayesian analysis indicated weak evidence in favour of the null hypothesis of no difference in imageability between the two categories of stimuli (BF_10_=.831). Thus, it is possible that the activity localised in bilateral motor cortices in the study by Vanhoutte et al. stems from a differential process of, potentially implicit, mental imagery between categories, that is absent from our data due to the more closely matched levels of imageability. Nevertheless, Vanhoutte et al. (2015), and others^62–64^, argue that the onset of this post-stimulus effect is too quick to reflect mental imagery. Conversely, critics of strong embodied accounts argue that such apparent rapid activation of motor cortices is not sufficient evidence for motor cortical involvement in representing the concept itself, as the activity could also reflect spreading activation from an abstracted representation^1^. It is also possible that other psycholinguistic variables that we did not measure, like for instance, how ‘arousing’ a word is to the reader^65^, differ between our word categories and drive our observed electrophysiological effects. Nevertheless, while our ERP data are consistent with differential semantic activations between verb categories, they do not provide evidence for a specific contribution of the cortical regions implicated by an embodied account.

Alongside an early ERP effect, we also observed a differential modulation of the mu rhythm (8-12Hz) in a late time-window (555-785ms). While the differences in power of the mu rhythm in this time-window did not exceed our statistical threshold at the sensor level (p=.050 where alpha=.025), a global dissimilarity analysis indicated strong evidence for different scalp distributions of the mu rhythm ERD (p=.009). As in the case of the ERPs above, this result suggests that the mu ERDs in response to motor and non-motor verbs are not generated by entirely overlapping regions of cortex. Furthermore, our source analyses provided strong evidence (BF_10_=10.598, p=.029 FDR-corrected) for a generator of this difference within the left superior parietal lobule – a region identified in a meta-analysis of fMRI and PET studies of lexical-semantic processing of action words or images^33^. Watson et al.^33^ suggested that the meta-analytical concordance in this parietal region stems from the use of object-directed action concepts across studies, as overlapping regions of parietal cortex are thought to support production of object-directed actions^66^, and lesions of parietal cortex are linked to deficits in recognising object-directed actions^67^. Indeed, many of the motor stimuli used in our study are concepts that describe object-directed actions – e.g. pull, hurl, carve.

Nevertheless, our mu data are also not consistent with an embodied view of action semantics as the effect is both late in time and not localised to specific sensorimotor cortices. The so-called mu rhythm (8-12Hz) in fact shares the same frequency band as the alpha rhythm but is differentiated by the fact that it is distributed over the top of the head with putative generators in rolandic regions^21^. However, our data did not show significant differences in this frequency band within rolandic brain regions. Indeed, our data provided substantial evidence for the null within left pre-central gyrus using Bayesian equivalent t-tests (BF_10_=0.236) – i.e. evidence that 8-12Hz power does not differ on average in this brain region during semantic access of both motor and non-motor verbs. It may therefore be more accurate to describe our observed 8-12Hz activity as reflecting the alpha rhythm, rather than the mu rhythm. The alpha rhythm has been characterised as an active inhibition mechanism whereby power increases functionally inhibit sensory processing in task-irrelevant brain regions, while power decreases boost processing in other regions^68^. In this sense, our observed greater decrease in 8-12Hz (alpha) power within the left superior parietal lobule in response to motor verbs may reflect a boost in processing in this brain region, perhaps as part of accessing the object-directed features of motor actions^66^. This finding is also in line with the stronger activation observed in fMRI and PET data in this region in response to action-related lexical stimuli^33^, as alpha power is often reported to be negatively correlated with the BOLD response of fMRI^69–71^.

Just as with the mu rhythm, beta band oscillations (13-30Hz) classically represent activation of the sensorimotor cortex upon imagining, executing, or observing a movement (e.g.^13–15^). However, we did not observe any effects in the beta band. Nevertheless, if we continue to assume that our observed 8-12Hz effect reflects the alpha rhythm rather than the mu rhythm, the lack of evidence for modulation of the beta band in our data is not surprising, and again fails to support an embodied view of motor action concepts.

While a number of authors report evidence for activity over motor regions during action-word processing and subsequently argue for an embodied account of action meaning, there are others who critically question this interpretation. For example, according to Bedny and Caramazza^72^, it is unclear whether the reported modulations over motor areas reflect the general role of these areas in language processing or reveals a specificity of motor areas in the understanding of action words. In a review article, the authors argue that there is more evidence for the role of left middle temporal gyrus in action word comprehension than sensorimotor regions – a position supported by the subsequent meta-analysis of Watson et al.^33^. Nevertheless, our findings at the source level do not provide evidence for a role of either of these cortical regions. As noted above, Bayesian t-tests of the alpha power effect provide evidence for the null hypothesis within the left pre-central gyrus, consistent with a more critical view of semantic embodiment of action verbs. Furthermore, our source analyses of the early ERP effect also revealed substantial evidence in favour of the null within the posterior middle temporal gyrus.

In conclusion, our data are consistent with a rapid differential activation of cortex when accessing the meaning of motor and non-motor verbs, followed by a later post-lexical involvement of left superior parietal lobule. However, our data do not provide direct support for a specific role of sensorimotor cortices when healthy individuals access the meaning of individual motor action verbs. To further delineate the extent to which embodied cognition applies to semantic representations, we must continue to delineate the specific task goals and/or contexts in which sensorimotor cortices are recruited in service of comprehension^34^.

## Author Contributions

SC and DC designed the task and stimuli. SC collected the data. SC, RS, and DC conducted the analyses. RS and DC wrote the first draft. All authors contributed to subsequent drafts.

## Additional Information

All authors declare no competing interests.

## Acknowledgments

This research was supported by generous funding from the Medical Research Council (MR/P013228/1). SC was supported by the Erasmus+ for Traineeships program.

## References

1. Mahon, B. Z. & Caramazza, A. A critical look at the embodied cognition hypothesis and a new proposal for grounding conceptual content. Journal of Physiology-Paris 102, 59–70 (2008).

2. Kiefer, M. & Pulvermüller, F. Conceptual representations in mind and brain: Theoretical developments, current evidence and future directions. Cortex 48, 805–825 (2012).

3. Glenberg, A. M. & Kaschak, M. P. Grounding language in action. Psychonomic Bulletin and Review 9, 558–565 (2002).

4. Glenberg, A. M., Sato, M. & Cattaneo, L. Use-induced motor plasticity affects the processing of abstract and concrete language. Current Biology 18, R290–R291 (2008).

5. Mirabella, G., Iaconelli, S., Spadacenta, S., Federico, P. & Gallese, V. Processing of Hand-Related Verbs Specifically Affects the Planning and Execution of Arm Reaching Movements. PLoS One 7, e35403 (2012).

6. Tettamanti, M. et al. Listening to Action-related Sentences Activates Fronto-parietal Motor Circuits. http://dx.doi.org/10.1162/0898929053124965 17, 273–281 (2006).

7. Aziz-Zadeh, L., Wilson, S. M., Rizzolatti, G. & Iacoboni, M. Congruent Embodied Representations for Visually Presented Actions and Linguistic Phrases Describing Actions. Current Biology 16, 1818–1823 (2006).

8. Raposo, A., Moss, H. E., Stamatakis, E. A. & Tyler, L. K. Modulation of motor and premotor cortices by actions, action words and action sentences. Neuropsychologia 47, 388–396 (2009).

9. Willems, R. M., Toni, I., Hagoort, P. & Casasanto, D. Neural Dissociations between Action Verb Understanding and Motor Imagery. Journal of Cognitive Neuroscience 22, 2387–2400 (2010).

10. Kemmerer, D., Castillo, J. G., Talavage, T., Patterson, S. & Wiley, C. Neuroanatomical distribution of five semantic components of verbs: Evidence from fMRI. Brain and Language 107, 16–43 (2008).

11. Hauk, O., Johnsrude, I. & Pulvermüller, F. Somatotopic Representation of Action Words in Human Motor and Premotor Cortex. Neuron Neuron 41, 301–307 (2004).

12. Pfurtscheller, G. & Lopes da Silva, F. H. Event-related EEG/MEG synchronization and desynchronization: basic principles. Clinical Neurophysiology 110, 1842–1857 (1999).

13. Cheyne, D. O. MEG studies of sensorimotor rhythms: A review. Experimental Neurology 245, 27–39 (2013).

14. Pfurtscheller, G., Brunner, C., Schlogl, A. & Lopes da Silva, F. H. Mu rhythm (de)synchronization and EEG single-trial classification of different motor imagery tasks. Neuroimage 31, 153–159 (2006).

15. Muthukumaraswamy, S. D. & Johnson, B. W. Changes in rolandic mu rhythm during observation of a precision grip. Psychophysiology 41, 152–156 (2004).

16. Pfurtscheller, G. Central Beta Rhythm During Sensorimotor Activities in Man. Electroencephalography and Clinical Neurophysiology 51, 253–264 (1981).

17. Pfurtscheller, G., Linortner, P., Winkler, R., Korisek, G. & Müller-Putz, G. Discrimination of Motor Imagery-Induced EEG Patterns in Patients with Complete Spinal Cord Injury. Computational Intelligence and Neuroscience 2009, 1–6 (2009).

18. Müller-Putz, G. R. et al. Event-related beta EEG-changes during passive and attempted foot movements in paraplegic patients. Brain Res. 1137, 84–91 (2007).

19. Pfurtscheller, G., Leeb, R., Keinrath, C. & Friedman, D. Walking from thought. Brain Res. (2006).

20. Cruse, D. et al. Detecting Awareness in the Vegetative State: Electroencephalographic Evidence for Attempted Movements to Command. PLoS One 7, e49933 (2012).

21. Ritter, P., Moosmann, M. & Villringer, A. Rolandic alpha and beta EEG rhythms’ strengths are inversely related to fMRI-BOLD signal in primary somatosensory and motor cortex. Hum. Brain Mapp. 30, 1168–1187 (2009).

22. Hauk, O. & Tschentscher, N. The Body of Evidence: What Can Neuroscience Tell Us about Embodied Semantics? Front. Psychol. 4, (2013).

23. Niccolai, V. et al. Grasping Hand Verbs: Oscillatory Beta and Alpha Correlates of Action-Word Processing. PLoS One 9, e108059 (2014).

24. Pulvermüller, F., Lutzenberger, W. & Preissl, H. Nouns and Verbs in the Intact Brain: Evidence from Event-related Potentials and High-frequency Cortical Responses. Cerebral Cortex 9, 497–506 (1999).

25. Preissl, H., Pulvermüller, F., Lutzenberger, W. & Birbaumer, N. Evoked potentials distinguish between nouns and verbs. Neuroscience Letters 197, 81–83 (1995).

26. Hauk, O. & Pulvermüller, F. Neurophysiological Distinction of Action Words. Hum. Brain Mapp. 21, 191–201 (2004).

27. Pulvermüller, F., Härle, M. & Hummel, F. Walking or Talking?: Behavioral and Neurophysiological Correlates of Action Verb Processing. Brain and Language 78, 143–168 (2001).

28. Shtyrov, Y., Hauk, O. & Pulvermüller, F. Distributed neuronal networks for encoding category-specific semantic information: the mismatch negativity to action words. Eur. J. Neurosci. 19, 1083–1092 (2004).

29. Pulvermüller, F., Shtyrov, Y. & Ilmoniemi, R. Brain Signatures of Meaning Access in Action Word Recognition. http://dx.doi.org/10.1162/0898929054021111 17, 884–892 (2006).

30. Maguire, M. J. et al. Electroencephalography theta differences between object nouns and action verbs when identifying semantic relations. Language, Cognition and Neuroscience 30, 673–683 (2014).

31. van Elk, M., van Schie, H. T., Zwaan, R. A. & Bekkering, H. The functional role of motor activation in language processing: Motor cortical oscillations support lexical-semantic retrieval. Neuroimage 50, 665–677 (2010).

32. Pulvermüller, F., Preissl, H., Lutzenberger, W. & Birbaumer, N. Brain Rhythms of Language: Nouns Versus Verbs. Eur. J. Neurosci. 8, 937–941 (1996).

33. Watson, C. E., Cardillo, E. R., Ianni, G. R. & Chatterjee, A. Action Concepts in the Brain: An Activation Likelihood Estimation Meta-analysis. http://dx.doi.org/10.1162/jocn_a_00401 25, 1191–1205 (2013).

34. Willems, R. M. & Francken, J. C. Embodied Cognition: Taking the Next Step. Front. Psychol. 3, (2012).

35. Vanhoutte, S. et al. Early lexico-semantic modulation of motor related areas during action and non-action verb processing. J. Neurolinguist. 34, 65–82 (2015).

36. Faul, F., Erdfelder, E., Lang, A.-G. & Buchner, A. G*Power 3: A flexible statistical power analysis program for the social, behavioral, and biomedical sciences. Behavior Research Methods 39, 175–191 (2007).

37. van Casteren, M. & Davis, M. H. Match: A program to assist in matching the conditions of factorial experiments. Behavioural Research Methods 39, 973–978 (2007).

38. Coltheart, M. The MRC psycholinguistic database. The Quarterly Journal of Experimental Psychology 33, 497–505 (2007).

39. A, K. BNC database and word frequency lists.

40. Davis, C. J. N-Watch: A program for deriving neighborhood size and other psycholinguistic statistics. Behavior Research Methods 37, 65–70 (2005).

41. Cortese, M. J. & Fugett, A. Imageability ratings for 3,000 monosyllabic words. Behavior Research Methods, Instruments, & Computers 36, 384–387 (2004).

42. Kuperman, V., Stadthagen-Gonzalez, H. & Brysbaert, M. Age-of-acquisition ratings for 30,000 English words. Behavior Research Methods 44, 978–990 (2012).

43. Love, J. et al. JASP (Version 0.7.1) [Computer Software].

44. Morey, R. D. & Rouder, J. N. BayesFactor (Version 0.9.11-3) [Computer Software].

45. Brainard, D. H. The Psychophysics Toolbox. Spatial vision 10, 433–436 (1997).

46. Delorme, A. & Makeig, S. EEGLAB: an open source toolbox for analysis of single-trial EEG dynamics including independent component analysis. J. Neurosci. Methods 134, 9–21 (2004).

47. Tadel, F., Baillet, S., Mosher, J. C., Pantazis, D. & Leahy, R. M. Brainstorm: A User-Friendly Application for MEG/EEG Analysis. Computational Intelligence and Neuroscience 2011, 1–13 (2011).

48. Oostenveld, R., Fries, P., Maris, E. & Schoffelen, J.-M. FieldTrip: Open Source Software for Advanced Analysis of MEG, EEG, and Invasive Electrophysiological Data. Computational Intelligence and Neuroscience 2011, 1–9 (2011).

49. Skrandies, W. Global field power and topographic similarity. Brain Topography 3, 137–141 (1990).

50. Maris, E. & Oostenveld, R. Nonparametric statistical testing of EEG- and MEG-data. J. Neurosci. Methods 164, 177–190 (2007).

51. Beukema, S. et al. A hierarchy of event-related potential markers of auditory processing in disorders of consciousness. NeuroImage: Clinical 12, 359–371 (2016).

52. Van Drongelen, W., Yuchtman, M., Van Veen, B. D. & van Huffelen, A. C. A spatial filtering technique to detect and localize multiple sources in the brain. Brain Topography 9, 39–49 (1996).

53. Van Veen, B. D., Van Drongelen, W., Yuchtman, M. & Suzuki, A. Localization of brain electrical activity via linearly constrained minimum variance spatial filtering. IEEE Trans. Biomed. Eng. 44, 867–880 (1997).

54. Robinson, S. Functional neuroimaging by synthetic aperture magnetometry (SAM). Recent advances in biomagnetism (1999).

55. Brookes, M. J. et al. A multi-layer network approach to MEG connectivity analysis. Neuroimage 132, 425–438 (2016).

56. Sokoliuk, R. et al. Two spatially distinct posterior alpha sources fulfill different functional roles in attention. bioRxiv 384065 (2018). doi:10.1101/384065

57. Benjamini, Y., B, Y. H. J. O. T. R. S. S. S.1995. Controlling the False Discovery Rate: A Practical and Powerful Approach to Multiple Testing on JSTOR. JSTOR doi:10.2307/2346101

58. Yekutieli, D. & Benjamini, Y. Resampling-based false discovery rate controlling multiple test procedures for correlated test statistics. Journal of Statistical Planning and Inference 82, 171–196 (1999).

59. Rouder, J. N., Speckman, P. L., Sun, D., Morey, R. D. & Iverson, G. Bayesian t tests for accepting and rejecting the null hypothesis. Psychonomic Bulletin and Review 16, 225–237 (2009).

60. Van Essen, D. C. et al. An integrated software suite for surface-based analyses of cerebral cortex. J Am Med Inform Assoc 8, 443–459 (2001).

61. Patterson, K., Nestor, P. J. & Rogers, T. T. Where do you know what you know? The representation of semantic knowledge in the human brain. Nat. Rev. Neurosci. 8, 976–987 (2007).

62. Gullick, M. M., Mitra, P. & Coch, D. Imagining the truth and the moon: An electrophysiological study of abstract and concrete word processing. Psychophysiology 50, 431–440 (2013).

63. Welcome, S. E., Paivio, A., McRae, K. & Joanisse, M. F. An electrophysiological study of task demands on concreteness effects: evidence for dual coding theory. Experimental Brain Research 212, 347–358 (2011).

64. West, W. C. & Holcomb, P. J. Imaginal, Semantic, and Surface-Level Processing of Concrete and Abstract Words: An Electrophysiological Investigation. http://dx.doi.org/10.1162/08989290051137558 12, 1024–1037 (2006).

65. Hauk, O., Davis, M. H. & Pulvermuller, F. Modulation of brain activity by multiple lexical and word form variables in visual word recognition: A parametric fMRI study. Neuroimage 42, 1185–1195 (2008).

66. Buxbaum, L. J., Kyle, K., Grossman, M. & Coslett, B. Left Inferior Parietal Representations for Skilled Hand-Object Interactions: Evidence from Stroke and Corticobasal Degeneration. Cortex 43, 411–423 (2007).

67. Buxbaum, L. J., Kyle, K. M. & Menon, R. On beyond mirror neurons: Internal representations subserving imitation and recognition of skilled object-related actions in humans. Cognitive Brain Research 25, 226–239 (2005).

68. Klimesch, W., Sauseng, P. & Hanslmayr, S. EEG alpha oscillations: The inhibition–timing hypothesis. Brain Research Reviews 53, 63–88 (2007).

69. Goldman, R. I., Stern, J. M., Engel, J., Jr & Cohen, M. S. Simultaneous EEG and fMRI of the alpha rhythm. Neuroreport 13, 2487–2492 (2002).

70. Laufs, H. et al. Where the BOLD signal goes when alpha EEG leaves. Neuroimage 31, 1408–1418 (2006).

71. Scheeringa, R. et al. Neuronal Dynamics Underlying High-and Low-Frequency EEG Oscillations Contribute Independently to the Human BOLD Signal. Neuron 69, 572–583 (2011).

72. Bedny, M. & Caramazza, A. Perception, action, and word meanings in the human brain: the case from action verbs. Annals of the New York Academy of Sciences 1224, 81–95 (2011).

