## Supplemental Material for "Dissociable electrophysiological correlates of semantic access of motor and non-motor concepts"

**Supplementary Materials**

Complete list of stimuli:

| MOTOR VERBS |
| --- |
| catch - skip - flick - rub - stir - scrub - push - grasp - pull - seize - chew - prod - smack - drag - shrug - squeeze - flex - throw - slam - snip - speak - pinch - bend - slap - hurl - hold - carve - reach - give - hit - grab - dab - stretch |
| **NON-MOTOR VERBS** |
| fail - pray - burn - grow - melt - shrink - wed - blush - teach - beg - praise - shine - plead - taint - cheat - faint - spoil - starve - count - fade - thaw - learn - boil - sweat - warn - scare - wilt - win - tease - join - swear - quit - tempt |
